# Osteogenic differentiation potential and marker gene expression of different porcine bone marrow mesenchymal stem cell subpopulations selected in different basal media

**DOI:** 10.1101/2020.04.27.063230

**Authors:** Sangeetha Kannan, Jyotirmoy Ghosh, Sujoy K. Dhara

**Affiliations:** Ph.D. scholar, Department of Biotechnology, Jain University, Bangalore-560011, Karnataka, India; Molecular Biology laboratory, ICAR-National Institute of Animal Nutrition and Physiology, Bangalore-560 030, Karnataka, India; Stem Cell Laboratory, Division of Animal Biotechnology, ICAR-Indian Veterinary Research Institute, Izatnagar, Uttar Pradesh-243122, India

**Keywords:** Porcine, bone marrow, basal media, mesenchymal stem cells, osteogenic differentiation

## Abstract

Multipotent porcine mesenchymal stem cells (pMSC) are indispensable for research and therapeutic use. Derivation and culture media might affect the selection of MSC subpopulation and thus the differentiation potential of cells. In this study we evaluated the effects of αMEM, aDMEM, M199, αMEM/M199, aDMEM/M199 and αMEM/aDMEM media on porcine bone marrow MSC derivation; pre-differentiation expression of ALP, COL1A1, SPP1 and BGLAP osteogenic marker genes at passage 5 and 10 pMSC; and differentiation potential of passage 5 pMSC. Morphological changes and matrix formation in osteogenic cells were evaluated by microscopical examination and calcium deposit in osteocytes was confirmed by Alizarin Red S staining. Results indicated media independent selection of different bone marrow MSC subpopulations with different surface marker gene expressions. Many pMSC subpopulations in different media had CD14+ expressing cells. We also observed basal media dependent changes in osteogenic markers expression and differentiation potential of pMSC. The αMEM/aDMEM media grown pMSC showed best osteogenic differentiation potential. We thus recommended the testing of αMEM/aDMEM mixed media in other species for pre-differentiation MSC culture that are intended for better osteogenic differentiation.

**Summary:** Pre-differentiation basal media influence osteogenic differentiation potential of mesenchymal stem cells (MSC). Among the tested media, αMEM/aDMEM was the best for pre-differentiation porcine MSC culture intending to use in osteogenesis.

## Introduction

Multipotent porcine mesenchymal stem cells (pMSC) are indispensable for research and therapeutic use. Worldwide there are about 1058 registered clinical trials testing the mesenchymal stem cell-based products for the treatment of ailments like musculoskeletal tissue injuries and degenerations, diabetes, multiple sclerosis, cardiovascular diseases, liver fibrosis etc (http://clinicaltrials.gov/). Bone marrow mesenchymal stem cell (MSC) is the most commonly used cells for such treatment. The advantages of MSC are they are plastic adherent hence are easy to culture. They are multipotent therefore, are differentiable into their own lineages, the osteocytes, adipocytes and chondrocytes; and trans-differentiable to myocytes (Wakitani et al., 1995), cardiomyocytes (Guo et al., 2018) and neuronal cells (Choong et al., 2007). Despite their remarkable potential in various applications, the major challenge in functional tissue regeneration is the difficulty in providing a proper environment to regulate their self-renewal and differentiation. The MSC have been described as cells relatively easy to expand in simple culture media, which is an advantage for their applications (Girdlestone et al., 2009). However, the expanded MSC population remains heterogeneous and contains subpopulations of morphologically and antigenically distinct cells (Phinney and Prockop, 2007). Very little is known about the subpopulations of MSC which are able to differentiate into cells of specific lineages. Also the culture conditions for selectively propagating the different fractions of bone marrow MSC have not been established (Szade et al., 2011).

The MSC has been used to treat the musculoskeletal tissue injuries by direct injection in the affected site (Connolly, 1998) and engrafting of bio-scaffold attached tissues (Zhou et al., 2016). In rabbit experimental model the MSC implant is found successful in repairing the damaged tendon (Young et al., 1998). Engrafting of bone marrow MSC is reported to improve the bone strength in treated children suffering from osteogenesis imperfect (Digirolamo et al., 1999), a group genetic disorders in which bones of the affected patient fracture/break easily, often from mild trauma or with no apparent cause.

During osteogenesis, pre-osteoblast cells are formed first from MSC which is a common precursor (progenitor) for both the osteogenic and chondrogenic cells (Nefussi et al., 1991). With the progress of differentiation the similar cells aggregates by a process called condensation that precede with the increase in number of committed pre-osteoblast cells (Hall and Miyake, 1995). The pre-osteoblast cells transform into osteoblasts which then secrete a non-mineralized matrix to form osteoid. Osteoid finally gives rise to mature osteocytes by mineralization (Franz-Odendaal et al., 2005).The entire process is regulated by the Runt-related transcription factor 2 (RUNX2) or the core-binding factor subunit alpha-1 (CBF-alpha-1) master regulator by controlling the expression of type 1 collagen A (COL1A1), alkaline phosphatase (ALP), bone gamma carboxyglutamate (gla) protein (BGLAP)/osteocalcin and osteopontin (SPP1) genes (Kirkham and Cartmell, 2007). It is shown that the transactivation of RUNX2 is dependent on the post-translational modification of the protein (Xiao et al., 1998) induced by bone morphogenic protein-2 (BMP-2) through p300 mediated acetylation (Jeon et al., 2006). Out of all the RUNX2 controlled genes, alkaline phosphatase (ALP) is a cell surface protein ubiquitously expressed by several cell types and is used as marker for screening preosteoblasts (Jaiswal et al., 1997). The ALP gets up-regulated with the progression of MSC osteogenesis within 2 days of induction (Golub and Boesze-Battaglia, 2007).The COL1A1 is an early marker of osteoblast. Increase of COL1A1 expression is observed during transformation of osteoprogenitor to pre-osteoblasts cells (Köllmer et al., 2013).The BGLAP is a small conserved non collagenous extracellular matrix protein expressed during late osteoblast differentiation (Weinreb et al., 1990). The BGLAP/osteocalcin is found abundantly expressed in the bone; the function is associated with the mineralization and matrix synthesis; and is considered as the late differentiation marker (Ducy and Karsenty, 1995).

Keeping the above background in mind and the fact that; basal media play an important role for proliferation, maintenance of undifferentiated states and differentiation potential of MSC (Brown et al., 2013), this study was designed to understand the effects of αMEM, aDMEM, M199, αMEM/M199, aDMEM/M199 and αMEM/aDMEM media on derivation, expression of RUNX2 dependant osteogenic marker genes viz., the ALP, COL1A1, SPP1 and BGLAP at passage 5 and 10, and the osteogenic differentiation potential of the porcine bone marrow mesenchymal stem cells isolated by different basal media at passage 5.

## Results

### Surface marker gene expression in different animals

The MSC derived from all the three animals had expressed the three positive markers the CD105, CD90 and CD73. The band intensity CD105 and CD73 although did not change, the CD90^+high^ and CD90^+low^ expression types were observed across the media in all the three animals. Among the negative markers the general leucocytes marker CD45 expression was absent in all except in aDMEM/M199 media. The expression of CD34 was low in most of the media however, no expression was observed in M199 in all the three animals. The CD14 expression was observed in all the animal bone marrow MSC derived in one or multiple basal media. The cells showed three different CD14^+high^, CD14^+low^ and CD14^-^ expression types in all the three animals (Fig. 1).

**Fig. 1.**
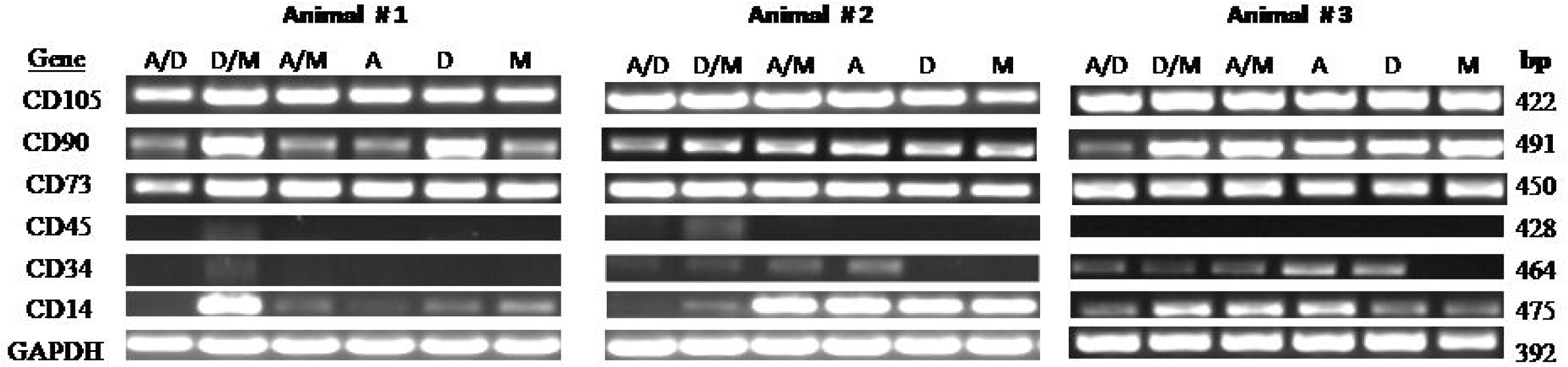
Surface marker gene expression of porcine bone-marrow mesenchymal stem cells derived from long bones of three different animals determined by PCR amplification after 40 cycles showing expression of positive markers: CD105 (endoglin), CD73 (SH3/4), CD90 (Thy-1) and negative markers: CD45 (leukocyte common antigen) CD34 (hematopoietic stem cell antigen) and CD14 (monocyte antigen) and endogenous control GAPDH (Glyceraldehyde 3-phosphate dehydrogenase) genes at passage 5 using gene-specific primers. Cells derived in six different basal media αMEM/aDMEM (A/D), aDMEM/M199 (D/M), αMEM/M199 (A/M), αMEM (A), advanced DMEM (D) and M199 (M) showed amplification of positive markers (CD105, Cd90 and CD73) in all the three animals however, amplification of CD90 was low in M, A, A/M, and A/D for animal # 1, M in animal #2 and A/D in animal #3. The leukocyte common antigen the CD45 expression was mostly not observed in all the 3 animals. Low level amplification of CD34 observed in D/M of animal #1, in A/D, D/M, A/M and A in animal #2 and in A/D, D/M, A/M, A and D of animal #3. Distinct CD14 expression was observed in D/M of animal #1; A/D, D/M, A/M and A of animal #2; and in D/M, A/M and A of animal #3. In other media CD14 amplification was either absent or low in all the three animals.

### Expression of osteogenic lineage marker gene expression in cells

All the media had shown ALP up-regulation in total and passage 5 cells. The up-regulation was also present at passage 10 cells of all media except in M199. At passage 10 compared to passage 5 the magnitude of up-regulation was reduced (p<0.05) in aDMEM, αMEM/aDMEM and aDMEM/M199 media but remained unchanged in M199 and αMEM/M199 (Fig. 2). The up-regulation was highest in αMEM/aDMEM than any other tested media. Overall the up-regulation of ALP (p < 0.05) was higher at passage 5 as compared to passage 10 (Fig. 2).

**Fig. 2.**
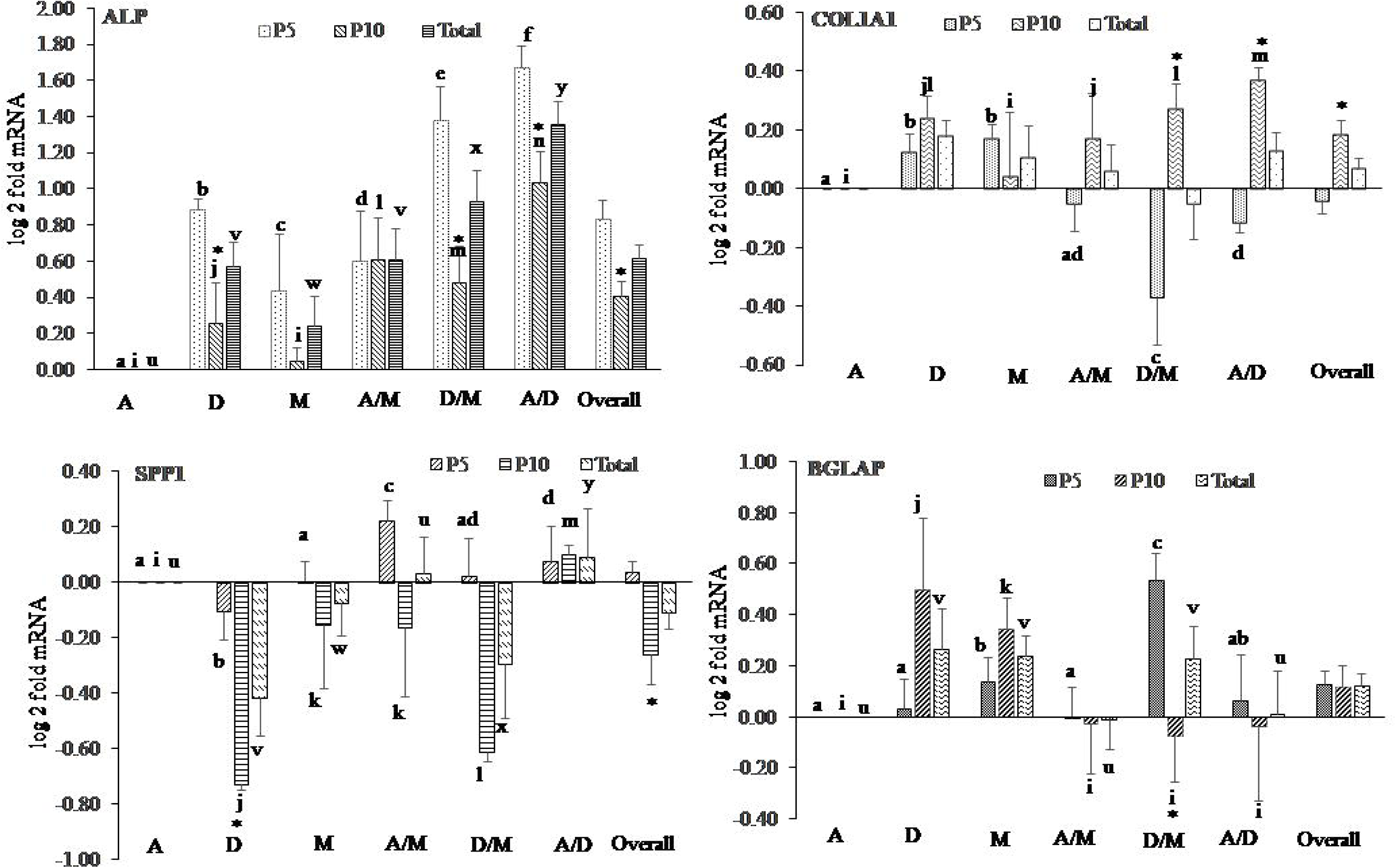
Media and passage wise effects on osteogenesis specific gene expression in porcine mesenchymal stem cells (pMSC). Relative mRNA expression of Alkaline phosphatase (ALP), Collagen type 1 (COL1A1), Secreted phosphoprotein1 (SPP1) and Bone gamma carboxyglutamate (gla) protein (BGLAP) in pMSC derived and grown in αMEM (A), aDMEM (D), M199 (M), αMEM/M199 (A/M), aDMEM/ M199 (D/M), αMEM/aDMEM (A/D) media at passage 5 (P5), passage 10 (P10) and overall. The GAPDH (Glyceraldehyde 3-phosphate dehydrogenase) was used as an endogenous control and the fold changes of gene expression were calculated using the method of Pfaffl, (2004) considering αMEM expression as calibrator. Values under each bar are mean ± SEM of triplicate samples from 3 animals. Different letters ‘a to f (P5), ‘i to n’ (P10) and ‘u to z’ (overall) indicated on top of each bar represented significant difference at p ≤ 0.05 between media. The ‘*’ on top of each bar represented significant difference at p ≤ 0.05 between the passages.

The total COL1A1 gene expression was up-regulated in αMEM/aDMEM compared to αMEM, and in aDMEM compared to aDMEM/M199. At passage 5 expressions was up-regulated in aDMEM and M199 and down regulated in aDMEM/M199 and αMEM/aDMEM media. At passage 10, up-regulation (p < 0.05) was observed in all the media except in M199 with the highest level in αMEM/aDMEM. Comparison between the passages revealed the down regulation in passage 5 was reversed to up regulation (p < 0.05) in passage 10 in aDMEM/M199 and αMEM/aDMEM media and in overall expression (Fig. 2).

The total SPP1 gene expression was down-regulated (p < 0.05) in aDMEM, M199 and aDMEM/M199 and up regulated in αMEM/aDMEM. At passage 5 however, this gene was down regulated in aDMEM and up regulated in αMEM/M199 and αMEM/aDMEM media. At passage 10 the media aDMEM, M199, αMEM/M199 and aDMEM/M199 shown down regulation however, the αMEM/aDMEM only maintained up-regulation (p < 0.05). Between the passages comparison revealed more (p < 0.05) down regulation of this gene overall and in aDMEM media at passage 10 than passage 5 (Fig. 2).

The total BGLAP expression was up-regulated in aDMEM, M199 and aDMEM/M199 media. At passage 5, the M199 and aDMEM/M199 media shown up regulation (p < 0.05) and at passage 10, the aDMEM and M199 media cells showed up-regulation. Comparison between the passages revealed significant reduction of expression in aDMEM/M199 at passage 10 as compared to the passage 5 level. However, the overall total expression of this gene did not change when the media effects were not considered (Fig. 2).

### Media interactions to affect the changes in gene expression

Interaction analysis revealed that all the three mixed media αMEM/M199, aDMEM/M199 and αMEM/aDMEM affected (p<0.05) the expression of SPP1. In addition to this gene the αMEM/M199 and aDMEM/M199 media also affected (p<0.05) the expression of COL1A1, and BGLAP genes and αMEM/aDMEM affected the expression of ALP (Fig. 2). The individual media αMEM only affected (p<0.05) the expression of SPP1 however, the aDMEM and M199 affected the expressions of the other three tested genes COL1A1, SPP1, and BGLAP (Fig. 3).

**Fig. 3.**
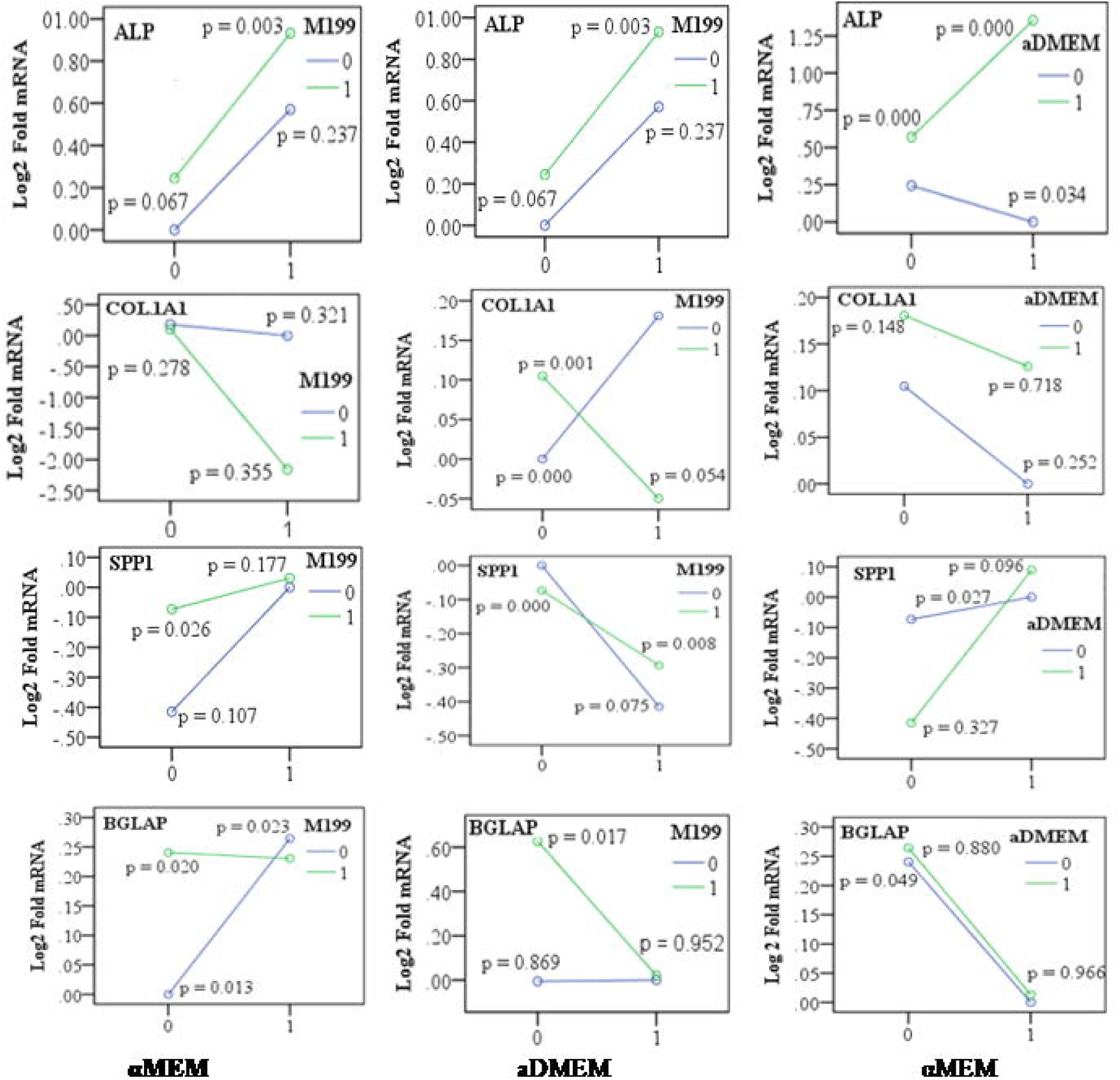
The media interaction for the changed expression of ALP, COL1A1, SPP1 and BGLAP in porcine mesenchymal stem cells (pMSC). The pMSC were derived and propagated in the absence (0, 0), presence of single (1, 0 / 0, 1) and combinations (1, 1) of two different basal media formulations. Gene expression in the absence of a particular media signified that the cells were cultured in a basal media other than the two particular media combinations. Triplicate of each sample was analyzed by established qPCR of each gene, GAPDH (Glyceraldehyde 3-phosphate dehydrogenase) was used as an endogenous control gene and αMEM expression as calibrator. The p values indicated in graphs were significant at p ≤ 0.05. The data of log 2 Fold mRNA expression at passage 5 and passage 10 of three different animals were pooled to obtain the marginal means for plotting the graph.

### Osteogenic differentiation of pMSC derived and grown in different media

During the entire 21 days osteogenic differentiation period, porcine MSC derived and grown in different media formulations underwent considerable changes in their morphology (Fig. 4 & 5). Varying differentiation ability was observed in different MSC subpopulations grown in different basal media as per the set evaluation criteria. The αMEM grown MSC had shown the whirling growth pattern of typical undifferentiated MSC by Day 6. However, by 12 days few growing osteoblast cells were visible in areas of cellular aggregates which matured by latter half of the differentiation period. In addition several adipocytes and undifferentiated cells were also seen by the end of the differentiation period. The aDMEM and M199 grown cells displayed changes in morphology from spindle-shaped to cuboidal by day 6 of induction which became more apparent by day 12 although a number of adipocytes and undifferentiated cells were observed in aDMEM by day 21. Most differentiated cells in this media showed a florid array of processes, a typical of early osteoblasts phenotype. In M199, cells in areas formed the dense aggregates of mature osteoblasts. By day 21, in both aDMEM and M199 nodular structures were visible due to coalescing of cells by mineral forming matrices over the multi-layers of cells and the differentiated adipocytes continued to be present.

**Fig. 4.**
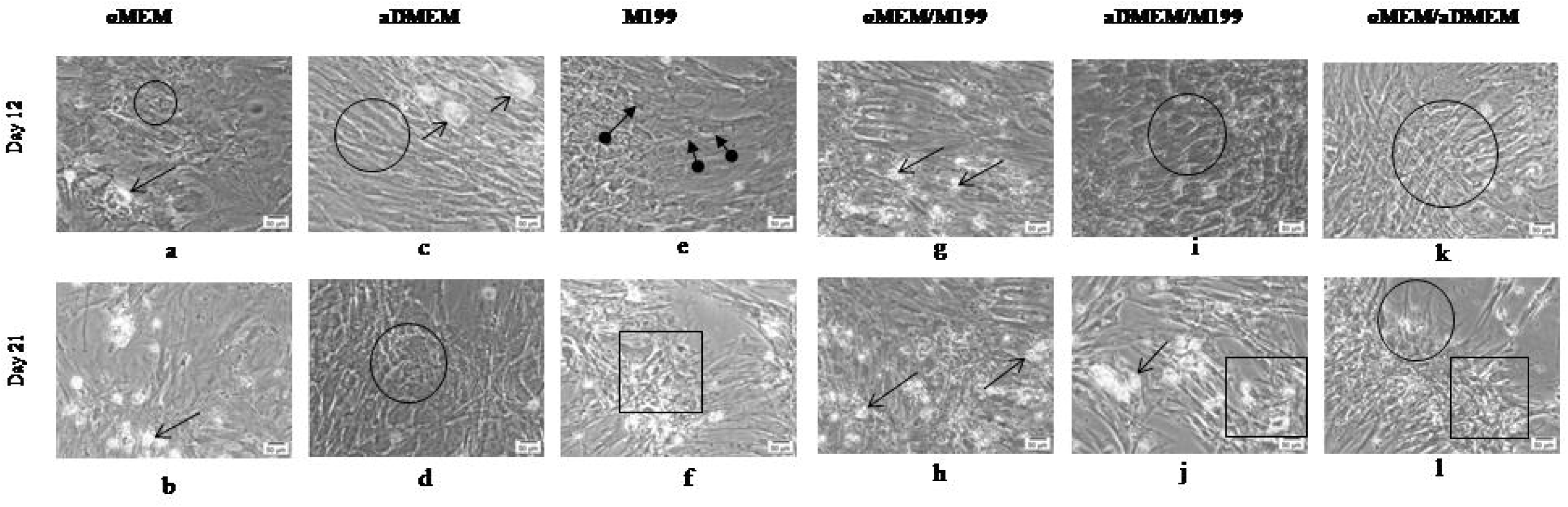
Phase contrast images showing a comparative morphology of porcine bone marrowderived mesenchymal stem cells on Day 12 and Day 21 of induction to osteogenic differentiation. The cells from αMEM formed very few osteoblasts (circle) at Day 12 (a) with also few adipogenic cells (opened arrow) as with αMEM/M199 (g) and aDMEM (c). Adipocytes were also present at Day 21 in αMEM, αMEM/M199 and aDMEM/M199 (b, h, j). Cells from aDMEM, aDMEM/M199 and αMEM/aDMEM displayed changes in their morphology to cuboidal shape typical of mature osteoblasts (circle) at Day 12 (c, i, k) and aDMEM and αMEM/aDMEM at Day 21 (d, l). In M199, a few cells tending towards osteogenesis (closed arrow) were seen at Day 12 (e) most of which matured by Day 21 (f). Cells in M199, aDMEM/M199 and αMEM/aDMEM formed nodular structures due to coalescing cells with visible mineral matrix (rectangular) by Day 21 of induction (f, j, l). Magnifications of respective images are as denoted by the scale bar.

In αMEM and αMEM/M199 media grown cells osteoblast cells were not detected during early period of induction (Fig. 5). However by day 12, a mixed population of osteoblasts, adipocytes and undifferentiated cells were seen which were continued to be present even after 21 days of differentiation and number of scanty osteocyte-like cells was observed in the canaliculi. Interestingly, cells of aDMEM/M199 and αMEM/aDMEM media showed morphological changes as early as day 3 of induction (Fig. 5) which progressed further by day 12 when most cells were cuboidal in shape (Fig. 4). In aDMEM/M199, a few adipocytes were also seen contaminating the osteoblast population. Although, distinct nodule aggregates exposing the bare plastic surface appeared in both the media, by day 21 number of contaminating adipocytes were present in aDMEM/M199 but not in the other media. Better calcified matrix depositions were visible in αMEM/aDMEM media with some cells transformed into stellar-shaped osteoids indicating osteoblast maturity. The pMSC grown in αMEM/aDMEM were bigger in sizes with granular cytoplasm an indication of skewing towards terminal differentiation (Fig. 5). During differentiation these cells displayed a well-developed extracellular matrix and multi-layered nodular structure throughout the culture dish with no contaminating adipocytes (Fig. 4). Extensive mineralization and calcium depositions was observed in these cells after 21 days of differentiation as evidenced by the Alizarin Red S staining which were not observed in other media grown cells (Fig. 5).

**Fig. 5.**
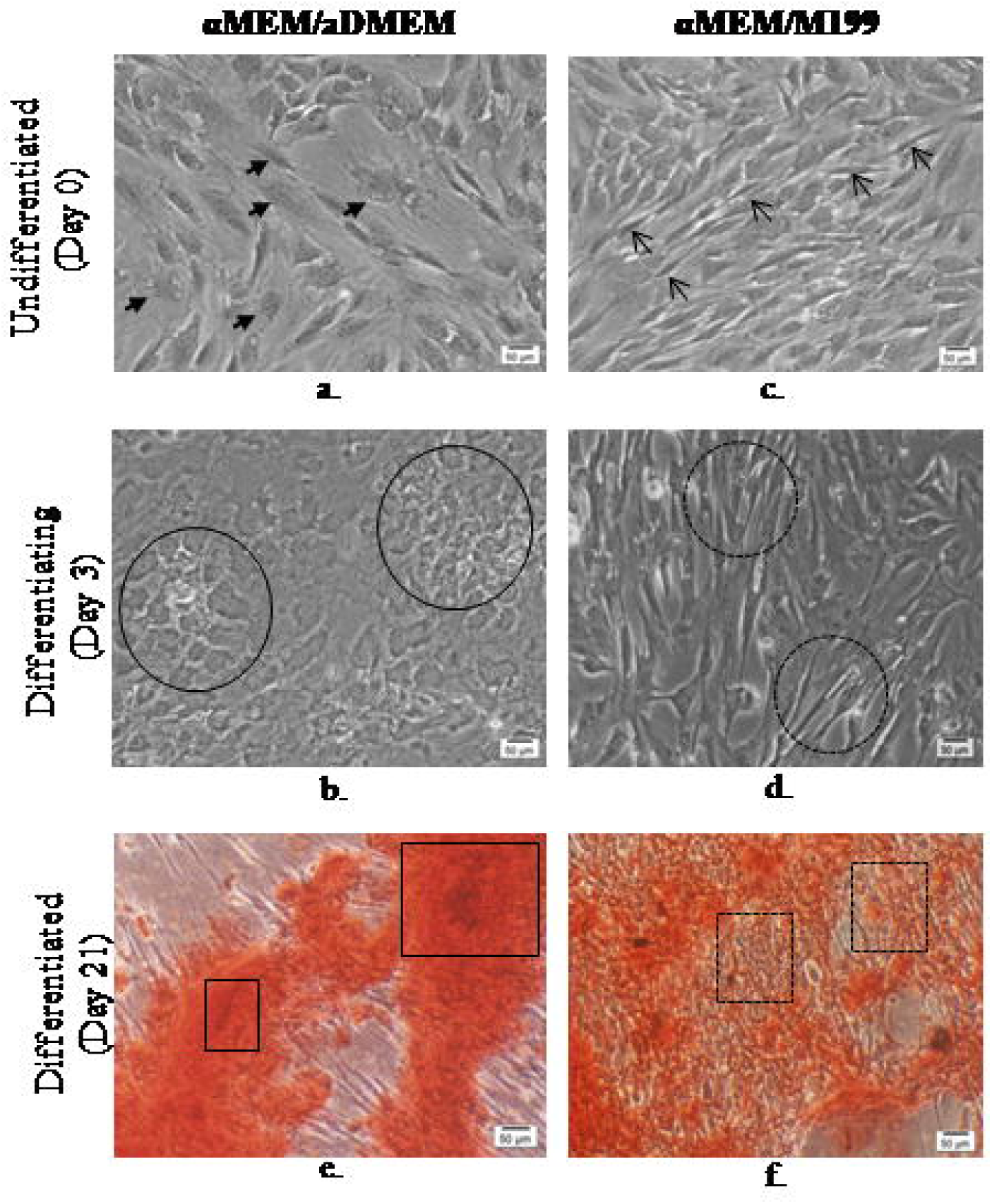
Phase-contrast micrographs of undifferentiated, osteogenically differentiating (day 3) and differentiated Passage 5 porcine bone marrow-derived mesenchymal stem cells. αMEM/aDMEM grown undifferentiated cells (a) showing larger cell size (closed arrows) compared to αMEM/M199 cells which were spindle-shaped (c) a typical of MSC morphology (opened arrows showing representative cells). Induction of osteogenic differentiation of αMEM/aDMEM grown cells (b) changed to cuboidal shape (line circles) as early as by day 3 of differentiation whereas cells grown in αMEM/M199 (d) did not show appreciable changes in morphology (dotted circles) on the same day. The calcification assessed by Alizarin red S stain at 21 days of osteogenic differentiation showed αMEM/aDMEM grown pMSC (e) displayed heavily stained Alizarin red S positive mineral matrix areas (line rectangles) whereas the pMSC grown in αMEM/M199 (f) showed patches of undifferentiated fibroblast-like cells (dotted rectangles) with very few cells displaying mineral deposition. Magnifications of respective images are as denoted by the scale bar.

## Discussion

It is of paramount important to understand if basal media plays any role in isolation of morphologically and antigenically distinct MSC subpopulations from bone marrow; and pre-differentiation MSC culture intending to use for a particular lineage specific cell differentiation. In this article we reported the media independent isolation of different MSC subpopulation expressing varying cell surface marker genes. Basal media played an important role in changing expression of osteogenic differentiation marker genes and skewing cells towards osteogenesis in pMSC. We also reported that the pMSC grown in αMEM/aDMEM mixed media was the best for osteogein differentiation irrespective of the cell surface gene expression as compared to the cells isolated and grown in other media.

The bigger sizes of pMSC in this media grown cells (Fig. 5) probably indicated the commitment towards differentiation. These cells had highest expression of the osteoprogenitor marker the ALP (Fig. 1), an indication of the presence of osteoprogenitor cells. Cells grown in this media also had shown earliest change in cell morphology by day 3 of differentiation induction, the best osteocyte maturing ability as evident by maximum calcium depositions and no visible undifferentiated MSC and contaminating adipocytes at the end of 21 days osteogenic differentiation (Fig. 4 & 5). Media composition might have played a role in the observed differences in osteogenic differentiation potential. The combined αMEM/aDMEM and αMEM media had similar number of amino acids and vitamins constituents however, varied in concentration (Supplementary table1). Many of the components were half the concentrations of αMEM. The ascorbic acid and L-Proline were two components that were present in αMEM/aDMEM mixed media @ 0.14 mM and 20 mg/L, exactly half the quantity of αMEM. Both the components at different levels are known to influence the collagen synthesis and osteogenic differentiation differently (Takamizawa et al., 2004).

The ALP is an early bone differentiation marker. The mutations of ALP cause severe bone malformations (Liu et al., 2018). Expression of ALP peaks highest in pre-osteoblasts formed from the osteoprogenitor cells (Jaiswal et al., 1997). Since the COL1A1 level was low (Fig 1) the cells were not progressed to pre-osteoblast cell types. Pre-osteoblasts are cuboidal cells that produce plenty of type 1 collagen and as the cells progress towards osteogenic commitment show elevated alkaline phosphatase and osteonectin levels (Franz-Odendaal et al., 2006). The pMSC grown in other media did not show change of morphology in the beginning of differentiation induction, however, the αMEM/aDMEM media grown cells become cuboidal as early as day 3 of induction (Fig. 5). Osteoblasts generated from pre-osteoblasts show a larger eccentric nucleus with nucleoli and prominent RER (rough endoplasmic reticulum) and Golgi areas. They express a suite of bone markers such as bone sialoprotein, osteocalcin, type 1 collagen. Upon mineralization, further changes such as reduced ER and Golgi correspond to a decreased protein synthesis and secretion (Franz-Odendaal et al., 2006). More ALP expression coinciding with the low expression of BGLAP (osteocalcin) in the pMSC and vice versa in different media (Fig. 1) indicated an inverse relationship between these two genes. A decline of ALP expression with the increase in level of BGLAP has been reported by others (Golub and Boesze-Battaglia, 2007). Cells with high ALP expression lack cell division and modulate the synthesis of osteocalcin (BGLAP) for directing differentiation of pre-osteoblasts into osteoblasts (Franz-Odendaal et al., 2006). The osteoblast secrete active bone matrix and typically display a cuboidal morphology (Franz-Odendaal et al., 2006). The varying timing in appearance of cuboidal cells indicated varying response of the different media grown cells towards osteogenic differentiation. High expression of alkaline phosphatase (ALP) and low to high level of COL1A1 in different media grown cells probably indicated varying progression of MSC towards osteogenic commitment. It has been reported that increase in expression of COL1A1 results in down-regulation of early osteoblast markers ALP with the increase in bone cell maturation (Malaval et al., 1994).The persistent up-regulation of ALP and down regulation of late osteogenic cell markers (e.g., COL1A1 and SPP1) until passage 10 (Fig. 1) in the different media grown porcine MSC probably indicated the maintenance of proliferation ability. The cells grown in αMEM/aDMEM indeed had shown less proliferability and lower expression of different cell proliferation marker gene (data under review).

Osteocalcin (BGLAP) is the late differentiation marker. The siRNA mediated knock-down of osteocalcin results in the down-regulation of RUNX2, alkaline phosphatase, osteonectin and type 1 collagen indicating the role of osteocalcin not only limited to the mineralization process but also in modulating the expression of transcription factors that control osteogenic differentiation (Tsao et al., 2017). Expression of osteoblast markers such as osteopontin (SPP1) and osteocalcin (BGLAP) increase concomitantly with the advancement in bone formation and mineralization (Malaval et al., 1994). Osteopontin (SPP1) is a secreted phosphoprotein whose central role is not fully characterized. In the bone matrix, osteopontin interact with integrin through its arginine-glycine-aspartate (RGD) motif and enable bone cells to adhere to the mineral matrix (Morinobu et al., 2003). Osteopontin-deficient mice showed a significant increase in the mineral depositions relative to the wild type counterparts without altering the expression of osteogenic markers indicating its minimal role in osteoblast development (Holm et al., 2014). The role of osteopontin is suggested in osteoclastic bone resorption rather than osteoblastic bone formation (Morinobu et al., 2003). Osteopontin (SPP1) up-regulation in αMEM/M199 and αMEM/aDMEM coinciding with the down-regulation of BGLAP; and increased up-regulation of BGLAP in most of the media coincided with down-regulation of SPP1. Earlier studies reported that the expression of BGLAP requires the phosphorylation of all the MAP kinases (ERK1/2, JNK1/2 and p38) while SPP1 gene expression requires only JNK1/2 and p38 phosphorylation (Kim et al., 2014. The readiness of M199, aDMEM, aDMEM/M199 and αMEM/aDMEM media grown cells for osteogenic differentiation was indicated by the high SPP1 and BGLAP gene expressions and was reflected by the appearance of pre-osteoblasts cells upon induction of osteogenic differentiation (Fig. 4).

The porcine bone marrow cells derived in different media were positive for the surface markers CD105, CD90 and CD73 so satisfied the properties of MSC as per the previous report (Kannan et al., 2019; Dominici et al., 2006).The differential expression patterns of positive CD90 marker, and the CD 45, CD 34 and CD14 across the media and animals could be associated with chance of MSC subpopulation selection during derivation and subsequent propagation. Any particular media did not found favouring selection of cells expressing a specific surface marker types. Considering the fact that bone marrow harbours a heterogeneous subpopulation of morphologically and antigenically distinct cell types (Phinney and Prockop, 2007) ioslation varying MSC subpopulation in different media is natural. The characterization of MSC based on surface and other markers thus should be mandatory considering adherence of a particular MSC subpopulation depends on their availability and chance. The MSC isolated from the same or different animals expressing different surface markers could be associated with changed differentiation potential and will further be helped by the media in which they were grown. It has been reported that reduced expression of CD105, CD90 or CD73 is associated with adipogenic differentiation, damage repair from bone fracture, or osteogenic differentiation through mechanical stimulation (Hung et al 2004; Ode et al 2011; Wiesmann et al., 2006). Our results indicated different CD90, ALP and COL1A1 expression in different media grown pMSC (Fig. 1) which might have contributed to the better osteogenic differentiation potential (Fig 4 & 5). The Thy-1 (CD90) is considered as a more effective biomarker for isolation of a highly osteogenic subpopulation of MSC for bone tissue engineering (Chung et al., 2013). The CD90 expression is associated with osteoprogenitor cells (Chen et al., 1999; Nakamura et al., 2010). Highest CD90 expression is reported in proliferating osteoprogenitor cells but the level decreases as the cells progress through the matrix maturation and mineralization stages (Chen et al., 1999). A decrease in CD90 expression coinciding with the increase in collagen-I and osteonectin protein expression in MSC is an indication of osteoblast-like cells (Wiesmann et al., 2006). The CD90-null mice gain weight at a faster rate, and ectopic over expression of CD90 block adipogenesis (Woeller et al., 2015). A down-regulation of CD90 through shRNA based knockdown is found favouring both the osteogenic and adipogenic differentiation (Moraes et al., 2016). Our study points to the fact that MSC differentiations towards a particular lineage need a delicate balance of CD90 and other osteogenic marker gene expression. The CD90 is a transient marker for early differentiation and it decreases concomitantly with the increase in collagen type 1 (COL1A1) and osteonectin (ON) expression (Wiesmann et al., 2006). The activation of COL1A1 is reported to promote the osteoblastic phenotype in human osteo-progenitor cells cultured in DMEM/Hams-F12 media (Ignatius et al., 2005). Induction of extracellular matrix (ECM) is a major activity of osteogenic cell types that occur due to the activity of type 1 collagen. Mutation in COL1A1 gene coding for the type 1 collagen protein showed reduced bone mineral density and increased osteoporotic vertebral fractures in humans (Grant et al., 1996). By blocking the interaction of integrin with type 1 collagen, osteocalcin and alkaline phosphatase expression was prevented (Xiao et al., 1998). At passage 5 up-regulation of COL1A1 in aDMEM and M199 probably indicated activation of osteogenesis process in the cells. However, in aDMEM/M199 and αMEM/aDMEM the activation was not apparent from the observed down regulation signal of COL1A1 in the cells (Fig. 2). The differentiation potential of MSC increases the levels of calcium deposition in passage 10 as compared to the cells of passages between 5 – 6 where alkaline phosphatase levels was also found lower (Wall et al., 2007). Report of others indicated that passage 15 porcine MSC grown in aDMEM retain only the adipogenic differentiation ability not the osteogenic potential (Vacanti et al., 2005). The retention of differentiation potential could be the function of the basal media in which cells were grown. As the αMEM media was used by Wall et al. (2007) whereas aDMEM media was used by Vacanti et al. (2005). This view is further supported by our observation of differences in osteogenic differentiation ability of the cells grown in different basal media (Fig. 4). The highest expression of alkaline phosphatase in αMEM/aDMEM with low COL1A1 (Fig. 2) could have favoured osteogenic differentiation as a result better calcification observed in the differentiated cells (Fig. 5).

Under certain favourable *in vitro* conditions, the MSC differentiate into their own lineages, (Moroni et al., 2013) and the cells of other lineages (Wu et al., 2013). We observed here the influence of pre-differentiation media on the changes in osteogenic differentiation potential of pMSC. The best osteogenic differentiation potential in terms of earliest appearance of ostegenic cells (Fig. 5), no contaminating adipocytes and better calcified cells at the end of differentiation was observed in αMEM/aDMEM media grown cell (Fig. 4 & 5) indicating better preparedness of these MSC toward osteogenic commitment. However, some media grown pMSC responded with appearance of adipocytes that continued to increase (not in huge number) during 21 days of osteogenic differentiation. Spontaneous generation of adipocytes during osteogenic differentiation might be the proof of adipogenic capacity of the cells (Braun et al., 2010). However, such adipocytes in the transplantation site may not be desirable when intended to use for particular therapeutic purpose such as osteogenesis.

Presence of CD14, the monocyte marker porcine bone marrow MSC (Fig. 1) is another interesting finding. According to International Society for Cellular Therapy, the MSC should be negative for CD14. However, 5% of the human bone marrow MSC is reported to have the cross reacting CD14 immuno epitopes (Pilz et al., 2011). Prominent CD14 band in pMSC subpopulation isolated in different media in all the three animals indicated the high CD14 positive cells subpopulation. Existence of CD14 positive MSC is supported by the fact that, circulating CD14 monocytes is a source of progenitors that exhibit mesenchymal cell differentiation ability (Kuwana et al., 2003); the bone marrow MSC in mouse are CD14+ (Szade et al., 2011); MSC derived from the peripheral blood and the adipose tissue of equine express CD14+ (Braun et al., 2010) and; the CD14 expression in adipose derived equine MSC is reported to wane with the increases in the passage number (McIntosh et al., 2006). Recently CD14 expression in MSCs is shown to plays an important role in regulating the LPS dependant TLR4 signalling pathway and dictates the differential activation through AKT, NF-kB and P38 signals (Jiang et al., 2019).

In summary, results of this study indicated that basal media composition may not have played any role in selection of MSC subpopulations from the bone marrow mass during derivation rather it was a chance of availability to adhere on the plastic surface. The porcine bone marrow might contain substantial CD14 expressing MSC. The osteogenic marker gene expression differentiation potential of pMSC was due to the compositional differences of pre-differentiation culture media. The αMEM/aDMEM media was better for skewing of porcine MSC towards osteogenic differentiation than other five tested basal media. We therefore recommended testing this media in other species for pre-differentiation MSC culture that are intended for better osteogenic differentiation.

## Materials and methods

All three basal media αMEM, aDMEM, M199 and osteogenic differentiation media HiOsteoXL™used for the study were purchased from HiMedia Laboratories, Mumbai, India. The mixed media αMEM/aDMEM, αMEM/M199, and aDMEM/M199 were prepared by mixing them at 1:1 ratio. The fetal bovine serum and media supplements β-mercaptoethanol, Glutamax and Antibiotic-Antimycotic solution were purchased from Gibco Life Sciences, USA and reagents from Sisco Research Laboratories (SRL), Mumbai, India.

If otherwise not stated the cells while derivation, passaging and differentiations were maintained at 37°C temperature, 5% CO_2,_ and > 85% relative humidity.

### Derivation and culture of porcine bone marrow MSC

The porcine MSC was derived from slaughter house collected long bone marrow of three pigs using the published protocol (Kannan et al., 2019). In brief, the bone marrow mass was collected in a sterile tube containing PBS (pH 7.2)-Acid-Citrate-Dextrose solution and 1% Antibiotic-Antimycotic solution. The cells from bone marrow were mechanically dispersed and divided equally into six parts and re-suspended in αMEM, aDMEM, M199, αMEM/M199, aDMEM/M199 and αMEM/aDMEM that contained 10% FBS, 0.1 % β-mercaptoethanol, 4 mM Glutamax and 1% Antibiotic-Antimycotic solution as supplements. The floating non-adherent cells were removed by media changes at every 48 h during derivation. The cells were passaged at 70 – 80% confluence by treating with 0.25% Trypsin-EDTA (# T4049, Sigma-Aldrich, USA). The cells were continuously grown until passage 10 however, used at intermediate passages for characterization and experimentation.

### Testing of cell marker gene expression

The MSC of all the three animals used for this experiment were tested for the presence or absence of CD105, CD90, CD73, CD45, CD34 and CD14 cell surface markers using GAPDH as endogenous control genes (Supplementary table 2) by PCR as per the published protocol (Sangeetha et al., 2019).

### Cell culture, total RNA isolation and cDNA synthesis

The passage 4 and 9 cells of each animal (n = 3) grown in all the six different media were seeded in triplicates at a density of 2 x 10^5^ cells/cm^2^ in sterile 6-well plates and grown until 70 – 80 % confluence with media changes at every 48 h. At termination of culture the spent media was discarded, adhered cells were rinsed with 1X PBS, placing the plate on - 20°C cold gel pack and treated with 0.4 mL/well TRIzol® LS reagent (#10296028,Ambion™, Life Technologies, USA). The TRIzol treated cell lysates from each well were harvested and stored at –80°C until use. Total RNA from the TRIzol treated samples was extracted following the manufacturer’s protocol. The total RNA yields and purity was determined by 260 and 280 nm absorbance using NanoDrop ND-2000 UV-Vis Spectrophotometer (Thermo Fischer Scientific, USA). The integrity of RNA in denatured samples was determined by 1% agarose gel electrophoresis in 0.2 mL 3-(N-morpholino) propanesulfonicacid (MOPS) buffer (pH 7.0). Single-stranded cDNA from 4µg total RNA was synthesized by reverse transcription (RT) using Revert Aid minus First Strand cDNA synthesis Kit (#K1621, Thermo Scientific) following manufacturer’s protocol.

### qPCR primers designing, synthesis and testing of efficiency

The primers used for the test ALP, COL1A1, SPP1, BGLAP and endogenous control GAPDH genes were designed (Table 1) using Primer3 web tool (http://bioinfo.ut.ee/primer3-0.4.0/primer3/) and synthesized from a local supplier (M/s Xcelris Labs limited, Ahmedabad, Gujarat, India). The amplification efficiency of all the gene primers were tested in 15 μL reaction mixtures that contained either 6.5 μL of cDNA template (50, 10, 2 and 0.4 ng total) for test and endogenous control genes in duplicate, nuclease-free water for non-template control (NTC) in triplicate and diluted total RNA (5 ng) for negative reverse transcriptase (RT) in duplicate; 0.5 μL each forward and reverse primers (5 pM) and 7.5 μL SYBR 2X master mix. The amplification efficiency of each pair of gene-specific primers was calculated using the formula E = [10^(−1/slope value)^-1]*100, where slope value is slope between average Ct and log quantity). The calculated primer efficiency of ALP, COL1A1, SPP1, BGLAP and GAPDH was 2.04, 1.92, 2.09, 2.02, and 1.93, respectively.

**Table 1:**
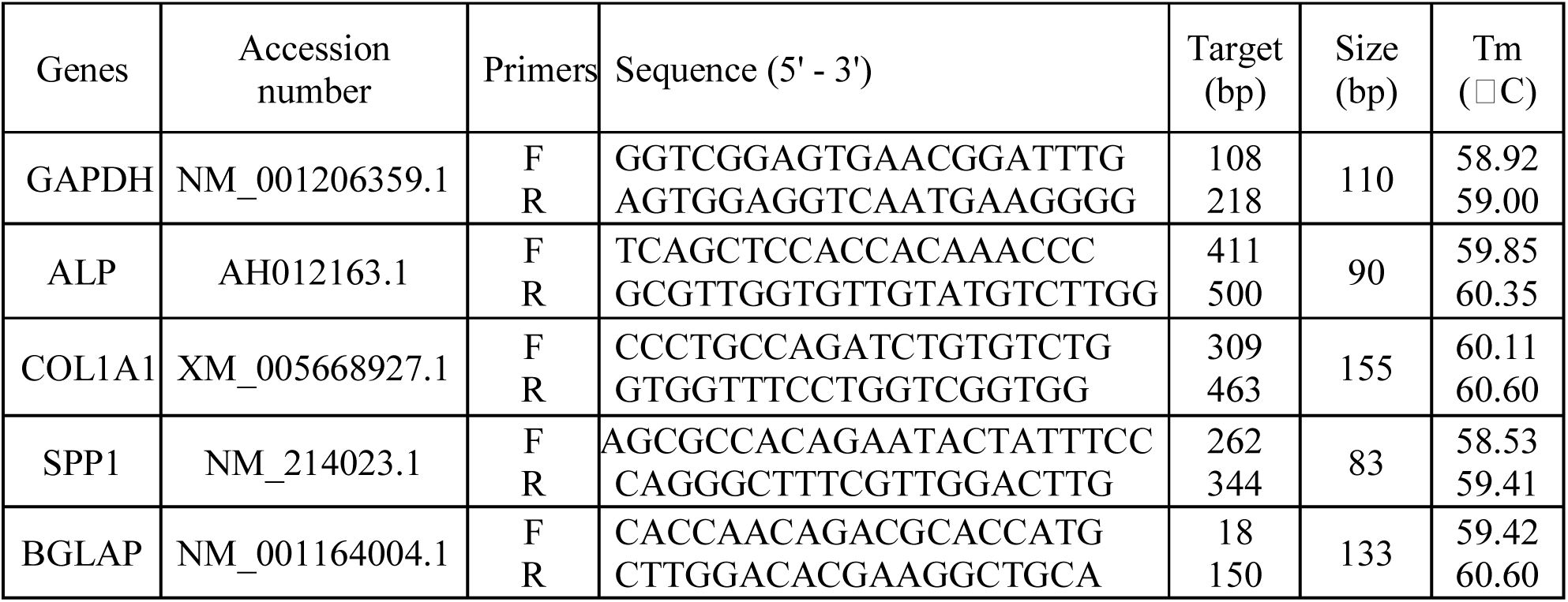
The list of qPCR primers used for relative quantification of osteogenic differentiation marker and endogenous control (GAPDH) genes with the accession number, sequence target (bp), amplicon sizes (bp) and melting temperature (Tm).

### qPCR analysis

The amplifications of target (ALP, COL1A1, SPP1 and BGLAP) and the endogenous control (GAPDH) genes were determined in 10 μL reaction mixtures of 96 wells clear Light Cycler 480 Multi well plate (#05102413001, Roche, USA). Triplicate 10 μL reaction mixtures were prepared that contained 4 μL either of 10 ng cDNA for gene of interest, 10 ng total RNA for negative reverse transcriptase control (-RT) and nuclease free water for non-template control (NTC), 0.5 μL of each forward and reverse primers (5 pM) and 5 μL of 2X FastStart SYBR Green master mix (# 6924204001, Roche, USA). The PCR cycling condition set in Light Cycler 480 instrument (Roche Diagnostics, USA)and was run at 95°C for 3 min initial denaturation followed by 50 cycles each comprising of 95°C denaturation for 30 s, 60°C annealing for 30 s and 72°C extensions for 30 s (image capture). The melt curve analysis was done at initial denaturation at 95°C for 1 min followed by annealing at 55 °C and increasing temperature 1°C per 10 s from 55 – 95°C and images captured at each 10 s intervals. The GAPDH was run for each plate and was used to normalize the variations in expression of target gene mRNA in each sample.

### qPCR analysis: Calculation of relative gene expression

The relative mRNA abundance of target genes in the sample was calculated from the comparative cycle threshold (Ct) using the formula (E_target_)^ΔCt_(target)_/(E_reference_)^ΔCt(_reference_) as described by Pffafl (2002). The ΔCt(_target_) was the difference between Ct of the target gene in calibrator (expression in αMEM) and target gene in the test (expression in other media) and ΔCt(_reference_) is the difference between Ct of the reference gene (GAPDH) in calibrator and reference gene in the test.

### Evaluation of cell morphology during osteogenic differentiation

The passage 4 cells of different media were seeded in 1 mL of respective media at a density of 5 x 10^3^ cells/cm^2^ per well in a sterile 24-well plate separately for all the animals (n=3). The media was changed with aDMEM containing all the supplements for 48 h uniformly and allowed to be grown until about 90 – 95 % confluence as per the subjective evaluation. The spent media was removed and replaced with equal volume of osteogenic differentiation medium in each of the wells. Fresh induction media were replaced at every 3 days until 21 days of differentiation. Changes in morphology and mineral matrix formation were recorded every day at 100X magnification using Olympus IX51 inverted microscope (Olympus, Japan). Cell images were captured at every 3 days using DP-73 camera and CellSens Entry 1.8 software (Olympus Soft Imaging Solutions, GmbH, Germany) and were evaluated for the comparative differences in morphology and progression of osteogenic differentiation based on the criteria for formation of mineral matrix or nodular structures, appearance of adipocytes and the undifferentiated cells remained in culture. The images were analysed by a single person from the sequential cell images captured at different time points.

### Confirmation of calcification in osteocytes by Alizarine red S staining

After 21 days of differentiation, the representative cells were stained using EZStain™ Osteocyte staining kit (Cat # CCK030; Himedia, Mumbai, India) to understand the deposition of mineral matrix in the differentiated cells. In brief, spent media from the wells were discarded and the cells were washed with PBS and incubated in ice-cold 70 % ethanol for 1 h for fixing. The ethanol from the cell surface was washed away by rinsing with sufficient water. The staining solution (pH 4.1-4.3) from the kit was added and incubated for 45 min at room temperature in dark. Finally the excess staining solution was removed by rinsing with distilled water. The stained cell images were captured by DP-73 camera using 200x magnifications in CellSens Entry 1.8 software (Olympus Soft Imaging Solutions, GmbH, Germany).

### Statistical analysis

The log 2 fold mRNA expression data of passage 5, passage 10 independently and pooled passage 5 and 10 were analyzed by full factorial design using General linear model and Multivariate analysis option. In which calculated Log 2 Fold gene expression was considered as dependent variables and media and passages as fixed factors. The significance of differences among the means was determined by LSD post hoc test of SPSS, 2010 Version 18.0 software. The interaction of two media combinations on different gene expression was analyzed by similar design, dependent variables and post hoc analysis but the two different media were used as fixed factors. The mean ±SEM values were considered significant in both the analysis at p ≤ 0.05.

## Acknowledgements

The Director ICAR-National Institute of Animal Nutrition and Physiology, Jain University, Bangalore and colleagues and supporting staff of Division of Physiology, ICAR-NIANP, Dr Debpriyo Dey, Senior Research Fellow of ICAR-All India Coordinated Research Project for the valuable inputs in interpretation of the images of cell differentiation.

## Author’s contributions

K.S. has performed the experiments, compiled the data, wrote the paper; J. G. conceived and designed the study, analyzed data, and wrote the paper; S. K. D. interpreted data and edited the manuscript.

## List of Symbols and Abbreviations used

αMEM: Minimum Essential Medium - alpha
aDMEM: Advanced Dulbecco’s Modified Eagle’s Medium
M199: Medium 199
PBS: Phosphate-Buffered Saline
EDTA: Ethylene diamine tetra acetate
MSC: Mesenchymal stem cells
pMSC: Porcine mesenchymal stem cells
RNA: Ribonucleic acid
mRNA: messenger Ribonucleic acid
shRNA: Short hairpin RNA
DNA: Deoxyribonucleic acids
UV: Ultraviolet
FBS: Fetal bovine serum
PCR: Polymerase chain reaction
qPCR: Quantitative polymerase chain reaction
Ct: Threshold cycle
RUNX2: Runt-related transcription factor 2
ALP: Alkaline Phosphatase
COL1A1: Type 1 collagen A1
SPP1: Secreted Phosphoprotein 1
BGLAP: Bone Gamma Carboxyglutamate (gla) Protein
GAPDH: Glyceraldehyde 3-phosphate dehydrogenase
AKT: RAC-alpha serine/threonine-protein kinase
NF-kB: Nuclear Factor kappa-light-chain-enhancer of activated B cells
JNK: c-Jun N-terminal kinase
BMP: Bone morphogenetic protein
LPS: Lipopolysaccharides
TLR: Toll-like receptors
p38: Protein 38
p300: Protein 300
SPSS: Statistical package for social sciences
SEM: Standard error of mean

